# Cell motion-coordinated fibrillar assembly of soluble collagen I to promote MDCK cell branching formation

**DOI:** 10.1101/869438

**Authors:** Jiajia Wang, Jia Guo, Bo Che, Mingxing Ouyang, Linhong Deng

## Abstract

Extracellular Matrix (ECM) assembly and remodeling are critical physiological events *in vivo*, and abnormal ECM assembly or remodeling is related to pathological conditions such as osteoarthritis, fibrosis, cancers, and genetic diseases. ECM assembly/remodeling driven by cells represents more physiological processes. Collagen I (COL) is very abundant in tissues, which assembly/remodeling is mediated by biochemical and mechanical factors. How cells regulate COL assembly biomechanically still remains to be well understood. Here we used fluorescent COL in the medium to study how cells assembled ECM which represents more physiological structures. The results showed that MDCK cells actively recruited COL from the medium and helped assemble the fibers, which in turn facilitated cell branching morphogenesis, both displaying high spatial associations and mutual dependency. Inhibition of cellular contraction force by ROCK and Myosin II inhibitors attenuated but did not block the COL fiber formation, while cell motion showed high consistency with the fiber assembly. Under ROCK or Myosin II inhibition, further analysis indicated high correlation between local cell movement and COL fiber strength as quantified from different regions of the same groups. Blocking cell motion by actin cytoskeleton disruption completely inhibited the fiber formation. These suggest that cell motion coordinated COL fiber assembly from the medium, possibly through generated strain on deposited COL to facilitate the fiber growth.

## Introduction

Epithelial branching (or tubular) structures exist broadly in organisms. Organs such as salivary glands, lungs, breasts and kidneys are composed of complex branching structures, and during embryonic development, the formation of these branching organs is called “branching morphogenesis” [1,2]. Mechanical force called biomechanics has important influence on developmental processes. Scientists even believe that mechanical and chemical factors are both indispensable for controlling embryonic development [3]. There have been studies on branching morphology on cell senses of physical factors in the environment, tension between cells and adjacent cells, and the effect of mechanical interaction between cells and extracellular matrix [4,5,6]. Recent study by Guo & Ouyang, et al. has found that ECM plays an important role in the formation of epithelial branching structure in MCF-10A cells, in which the long-range force between cells induces COL rearrangement and forms a mechanical feedback effect [7]. In the latest research, Baker et al. designed synthetic fibers with controllable stiffness, and found that cell stretching and network deformation led to recruitment of a large number of fibers around cells [8].

ECM assembly and remodeling are critical physiological events *in vivo*, and abnormal ECM assembly or remodeling is involved into pathological conditions such as osteoarthritis, fibrosis, cancers, and genetic diseases [9]. ECM assembly is mediated by both biochemical processes and mechanical force, and force on ECM regulates ECM remodeling, ligand binding, and homeostasis [10]. For instances, strain on fibronectin exposes the self-association sites to promote self-assembly [11,12,13]; fibre stretch-assay studies revealed that mechanical force regulates the interactions of fibronectin and Col [14]. Tension is required for fibrillogenesis of COL secreted by embryonic tendon cells [15]. Enzymatic degradation of COL fibrils is down-regulated by increasing mechanical stretch loading [16,17]. Basically, physical force is actively involved into ECM assembly and remodeling [18]. The mechanism of how cells regulate COL assembly from the medium biomechanically remains to be well understood.

In three-dimensional cell culture studies, COL often exists as extracellular matrix within the hydrogel. *In vivo*, the initial COL is secreted by the cells, and participates in the formation of collagen bundles [19]. COL assembly and remodeling driven cells represent more physiological processes. How COL molecules in medium form cells-assembled fibrillar structure remains unclear. In this work by using CNA35-EGFP-labeled COL in the medium, we visualized the developing process of cells-assembled COL fibers, and found cell contraction force-dependent motions regulated the fiber assembly from COL in the medium.

## Materials and Methods

### Cell culture and reagents

MDCK cells were purchased from the Beijing Beina Science & technology Co., Ltd. and cultured in Dulbecco’s Modified Eagle’s Medium (DMEM, high glucose; Thermo scientific) supplemented with 10% bovine fetal serum (FBS; Thermo scientific) and 1% antibiotic (penicillin & streptomycin; Thermo scientific). Cells were incubated at 37°C in a humidified incubator with 5% CO_2_.

Matrigel was purchased from BD Biotechnology, and type I collagen from rat tails from Advanced Biomatrix. Cytochalasin D, Blebbistatin, and Y27632 were purchased from Sigma. Plasmid pET28a-EGFP-CNA35 was purchased from Addgene [20]. Mouse monoclonal anti-collagen type I antibody (Clone COL-1), and Rhodamine-conjugated goat anti-mouse IgG antibody were purchased from Sigma.

### Cell culture on 3-D Matrigel

Firstly, we prepared polydimethylsiloxane (PDMS) (DOW) wells for making 3-D Matrigel. The E600 gel (Shenzhen Hongyejie Technology Co., Ltd) was poured into a 48-well plate, and formed individual cylinders with a diameter of 1 cm. The PDMS base and the cross-linking agent were thoroughly mixed at a mass ratio of 10:1, poured into a large dish with uniform insertion of the previously prepared cylinders, and then cured at 70 °C after vacuuming. After removal of the cylinders, PDMS wells were generated with a diameter of 1 cm and thickness of 0.5 cm. To assemble the PDMS molds, the prepared PDMS wells were plasma treated to achieve a more hydrophilic surface and then attached onto glass slides (Thermo Fisher Scientific).

To prepare the 3-D Matrigel, about 70 μl of Matrigel solution (BD Biotechnology) was added into a PDMS mold on ice, and then gelled at 37 °C for 30 min. COL molecules in solution (4 mg/ml) were labeled with EGFP-CNA35 protein (∼0.48 mg/ml) at volume ratio of 1:5 (EGFP-CNA35: COL) on ice for 10 min. Culture medium containing 1.5×10^4^ cells/cm^2^ and 20 μg/ml labeled COL was added on top of the 3-D Matrigel, and the samples were incubated in the cell culture incubator or imaged under live-cell microscopy.

Plasmid pET28a-EGFP-CNA35 was transformed into E. coli to produce EGFP-CNA35 protein. The expression and purification procedures were described in our previous work [21,22].

### Immunofluorescence

The samples cultured on 3-D Matrigel were first washed with PBS, fixed with 4% paraformaldehyde for 15 minutes, permeablized with 0.3% Triton-X 100 for 15 minutes, and blocked with 1% bovine serum albumin V (BSA) (BioFroxx, Germany). The immune-reaction was carried out overnight at 4 °C with the primary antibody. Mouse monoclonal anti-collagen type I antibody (Clone COL-1), and Rhodamine-conjugated goat anti-mouse IgG antibody were used at dilution of 1:500 and 1:100, respectively. The primary antibody was washed away with PBS, and the secondary antibody was added for staining at 37 ° C for 1 hour. Finally, the nucleus was stained with DAPI, and washed with PBS two minutes later. The fluorescence images were captured using the microscopy imaging system Zeiss cell observer (Karl Zeiss, Germany). Most of the imaging experiments on 3-D Matrigel were performed under a 20x objective.

### Mean fluorescence intensity quantification of COL fibers

A sample below displays the process to quantify the mean fluorescence intensity of COL fibers on the images. Briefly, in the software MatLab, the fluorescence on the image at the 24-hour time point (time 24) was subtracted by the same position at the start time point (time 0) as background. The resulted image (time 24-0) was switched into the format of gray (time 24-0 gray) by binary processing, in which the pixel values fell into the scale of 0-255. The mean value was calibrated based on the average value among those pixels with a threshold value above 10.

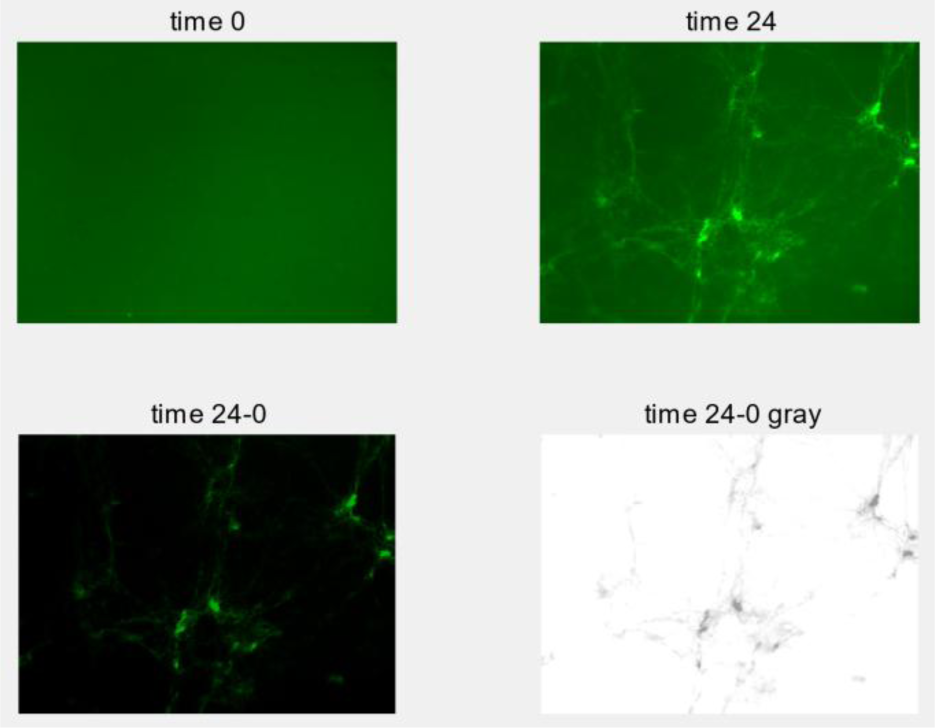

### Cell motion tracking

The image stack from the time course was processed by binarization in the software ImageJ. The positions of cell nuclei from a single cell were tracked along with the stack by using the function ‘manual tracking’ (Plugins -> Tracking -> Manual Tracking). The resulted readouts of coordinate (x, y) value from each time point were recorded into Excel (Microsoft) file, and further input into MatLab for calculation of move distance, and for graphic output of the migration trajectory.

### Statistical analysis

The values from different group data were plotted using GraphPad Prism6 software. Each value on the bar graphs represents the mean ± S.D (standard deviation). Student’s t-test was applied for statistical analysis, and P value < 0.05 was considered to be significant difference.

## Results

### Cells-dependent COL fiber assembly

In order to visualize COL fiber formation, COL molecules were labeled with fluorescent CNA35-EGFP, a COL binding protein [20,23]. MDCK cells (∼2×10^4^ cells/cm^2^) were cultured on 3-D Matrigel with 20 µg/ml fluorescence-labeled COL in the medium. In 24 h, MDCK cells formed branching structure along with fluorescent COL fibers on the Matrigel surface (Fig. 1A). In contrast, cells formed individual clusters and no fluorescent fibers were visible when culturing with CNA35-EGFP only without COL in the medium (Fig. 1B).

**Figure 1.**
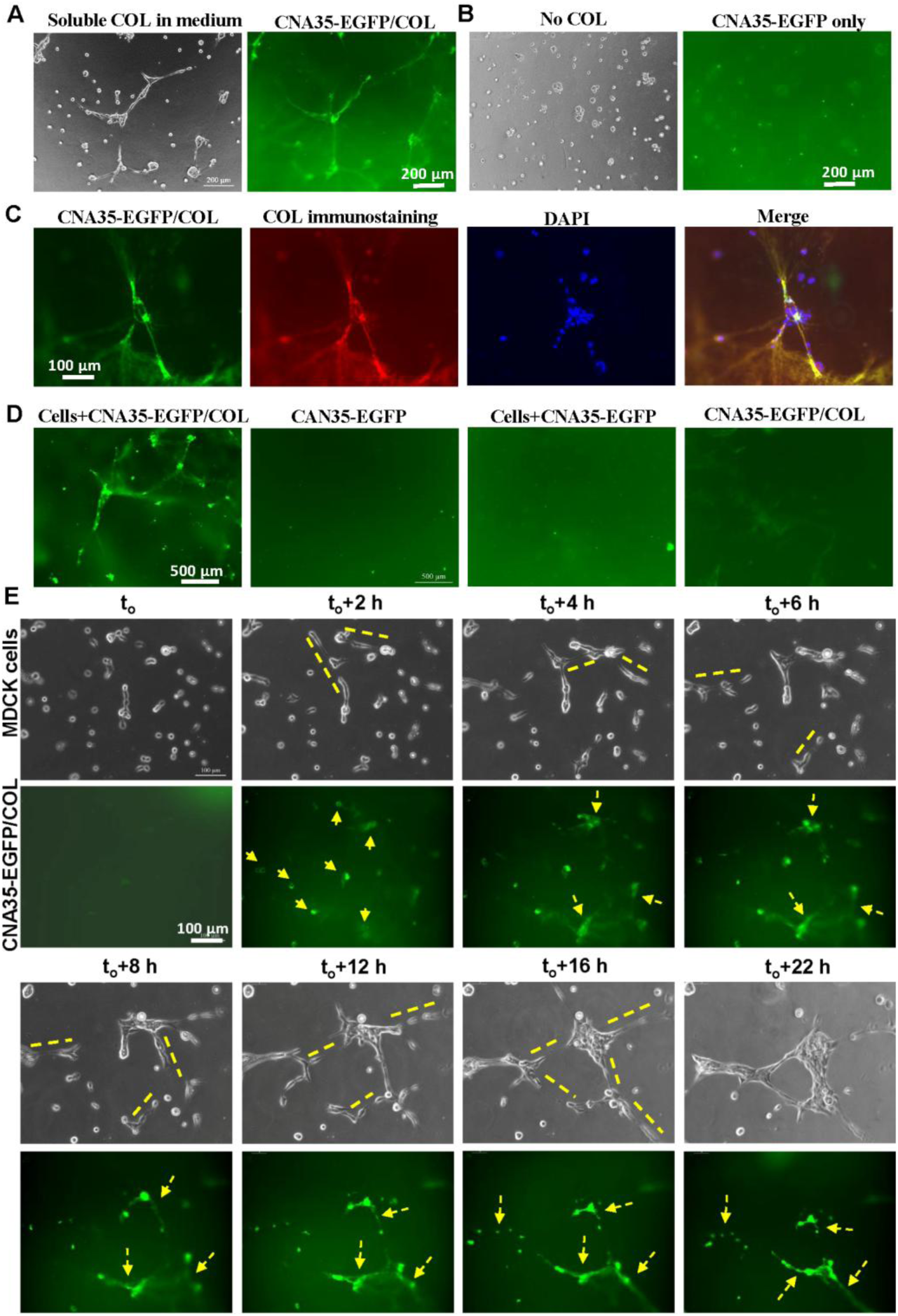
Cells-mediated COL fiber assembly from soluble molecules. MDCK cells (∼2×10^4^ cells/cm^2^) were cultured on 3-D Matrigel with 20 µg/ml fluorescence-labeled COL in the medium. (**A**) The formations of cell branching and fluorescent COL fibers in 24 h. (**B**) Cells formed clusters when with CNA35-EGFP but no COL in the medium. (**C**) Co-localization of CNA35-EGFP and immunostained COL with antibody. Cell nuclei were labeled with DAPI. (**D**) Check of COL fiber formation under the different culture conditions. (**E**) Simultaneous visualization of cell branching and COL fiber formations. MDCK cells were seeded on 3-D Matrigel with 20 ug/ml COL labeled with CNA35-EGFP in the medium. The time-lapse imaging with 30 min interval recorded the dynamic process of branching and fiber assembly (aslo seen in Movie S1). The lines on the images indicate the cell branching occurrence, and the arrows point to fluorescent COL fiber assembly.

To further confirm the fluorescent COL fibers, immunostaining of the samples with COL antibody (red) did display the co-localization with CNA35-EGFP (green) (Fig. 1C). The fluorescent fibers only showed up when co-existence of cells and CNA35-EGFP-labeled COL in the culture system, but were not obvious when containing CNA35-EGFP only, cells with CNA35-EGFP, or CNA35-EGFP-labeled COL only (Fig. 1D). These data together indicate the mutual dependence that cell branching formation required the existence of COL, and the fiber assembly from soluble COL also depended on cells.

### The dynamics of COL fiber assembly along with cell branching

The fluorescence-labeled COL allowed to check the dynamics how soluble COL in the medium was assembled into fibrillar structure. We used time-lapse imaging to simultaneously visualize cell branching formation and COL fiber assembly. After cells were seeded on the 3-D Matrigel, the dynamics was recorded as follows (shown in Fig. 1E and Movie S1): 1) cells started to initiate mutual connections for branching formation (indicated by the lines), and bright fluorescent spots appeared around cells in 2 h (pointed by the arrows), suggesting the possible recruitments of COL from medium by cells; 2) in 4-6 h, the fluorescent spots grew larger, and fibrillar collections between them started to show up, along with the appearance of cell branching structures at early stage; 3) in 8-12 h, the fibrillar structures got more apparent, the cell branching extended along with more cell-cell connections, and the two were associated spatially; 4) in 16-22 h, the COL fibers were assembled with sharp shape, and the cell branching was formed with grown strength, which two showed well spatial association. It was also seen that COL fibers were not uniformly distributed along with the cell branching, which might be partially due to lacking soluble COL later in the medium. From the dynamic process, the occurences of the cell branching and COL fiber assembly showed mutual dependence with close spatiotemporal associations.

### Cell contraction force-mediated COL fiber assembly

To further check whether cell contraction force regulates the mutual-dependent assembly of cell branching and COL fibers, we added ROCK signaling inhibitor Y27632 (40 µM) or Myosin II inhibitor Blebbistatin (25 µM) into the cell culture, which both inhibited cell contraction force. In comparison, cells formed longer branchings at the control condition (DMSO), whereas cells formed short branchings with either inhibitor in 24 hours (Fig. 2A(i-iii)). This was further confirmed by quantification of the branching length (Fig. 2B). COL fibers were still assembled with or without inhibitors, however, the fibers looked much stronger and sharper in morphology under control condition (Fig. 2A(i-iii)), which average of fluorescence intensity showed ∼60% stronger than those with inhibitors (Fig. 2C). The dynamic time courses of cell branching morphogenesis and fiber assembly were further displayed in Movies S2-4 (under control condition, or with Y27632 or Blebbistatin treatment). These data indicate certain dependence of the fiber assembly on cell contraction force. When addition of Cytochalasin D (Cyto D) (1 µM) to inhibit cell actin cytoskeleton, neither cell branchings nor COL fibers were assembled (Fig. 2A(iv), Movie S5), suggesting that the intact cytoskeleton was required for assembly of cell branching and COL fibers.

**Figure 2.**
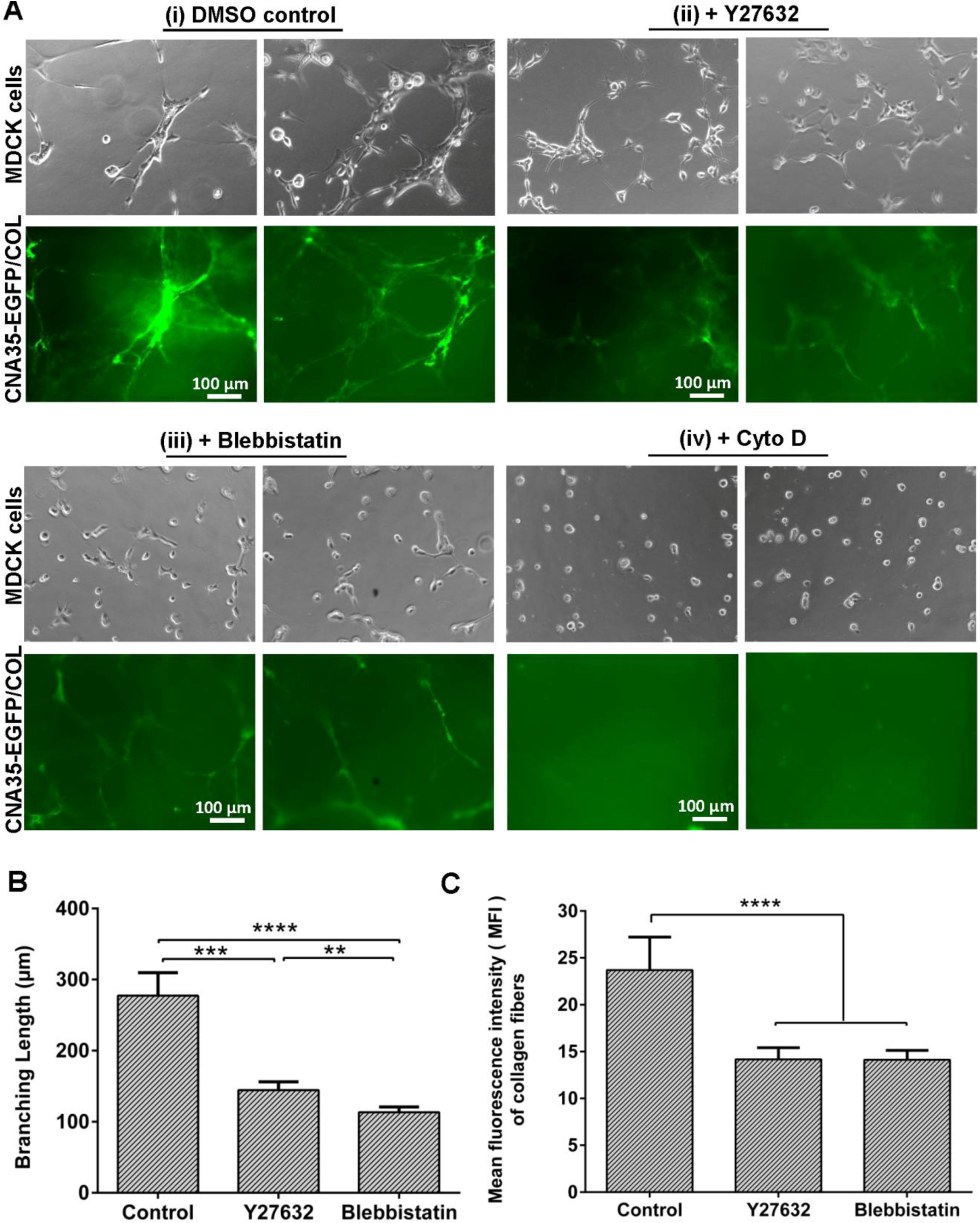
Cell contraction force-mediated assembly of cell branching and COL fibers. MDCK cells were cultured on 3-D Matrigel with soluble CNA35-EGFP/COL (20 µg/ml) in the medium containing control DMSO, Y27632 (40 µM), Blebbistatin (25 µM), or Cyto D (1 µM). Images were recorded with 30 min interval under microscopy for 24 h. (**A**) The assembly of cell branching and fluorescent COL fibers under control condition (DMSO) (i), or with the inhibitor Y27632 (ii), Blebbistatin (iii), or Cyto D (iv) in the culture medium (also seen in Movies S2-5). (**B**) Quantification of the branching length under control (DMSO), or with Y27632 or Blebbistatin. The lengths of the cell branching were measured and counted from multiple fields of microscopic views under 10x objective by using ImageJ software. No branching was formed with Cyto D treatment. (**C**) Quantification of the mean fluorescence intensity of assembled COL fibers under control (DMSO), or with Y27632 or Blebbistatin treatment. In statistical quantification, ** represents P < 0.01, *** P < 0.001, and **** P < 0.0001 for significant difference by Student’s t-test analysis.

### COL fiber assembly relying on cell motion

To further understand why inhibition of cell contraction force disturbed branching and COL fibers formations, we did quantification to track cell motions during the 24 hours seeding on the 3-D Matrigel. Under control condition, cells migrated in a more persistently directional way (Fig. 3A, Movie S2), whereas with Y27632 and Blebbitatin inhibitors, cells migrated more randomly without persistent directions (Fig. 3B-D). Inhibition of cell contraction force significantly attenuated cell mobility, as shown by the difference in total travel distance among the few conditions (Fig. 3E). Inhibition of Myosin II by Blebbistatin seemed to have more impact on cell motility than ROCK signaling by Y27632 (Fig. 3E). Quantification of final displacements showed that cells reached much smaller translocations at spatial scale under inhibition of cell contraction force than those under control condition (Fig. 3F), indicating inefficient directional migration after contraction force inhibition. Inhibition of intact actin cytoskeleton almost blocked cell translocation (Fig. 3F), which was consistent with the observation that no cell branchings or COL fibers were assembled under Cyto D treatment (Fig. 2A(iv)).

**Figure 3.**
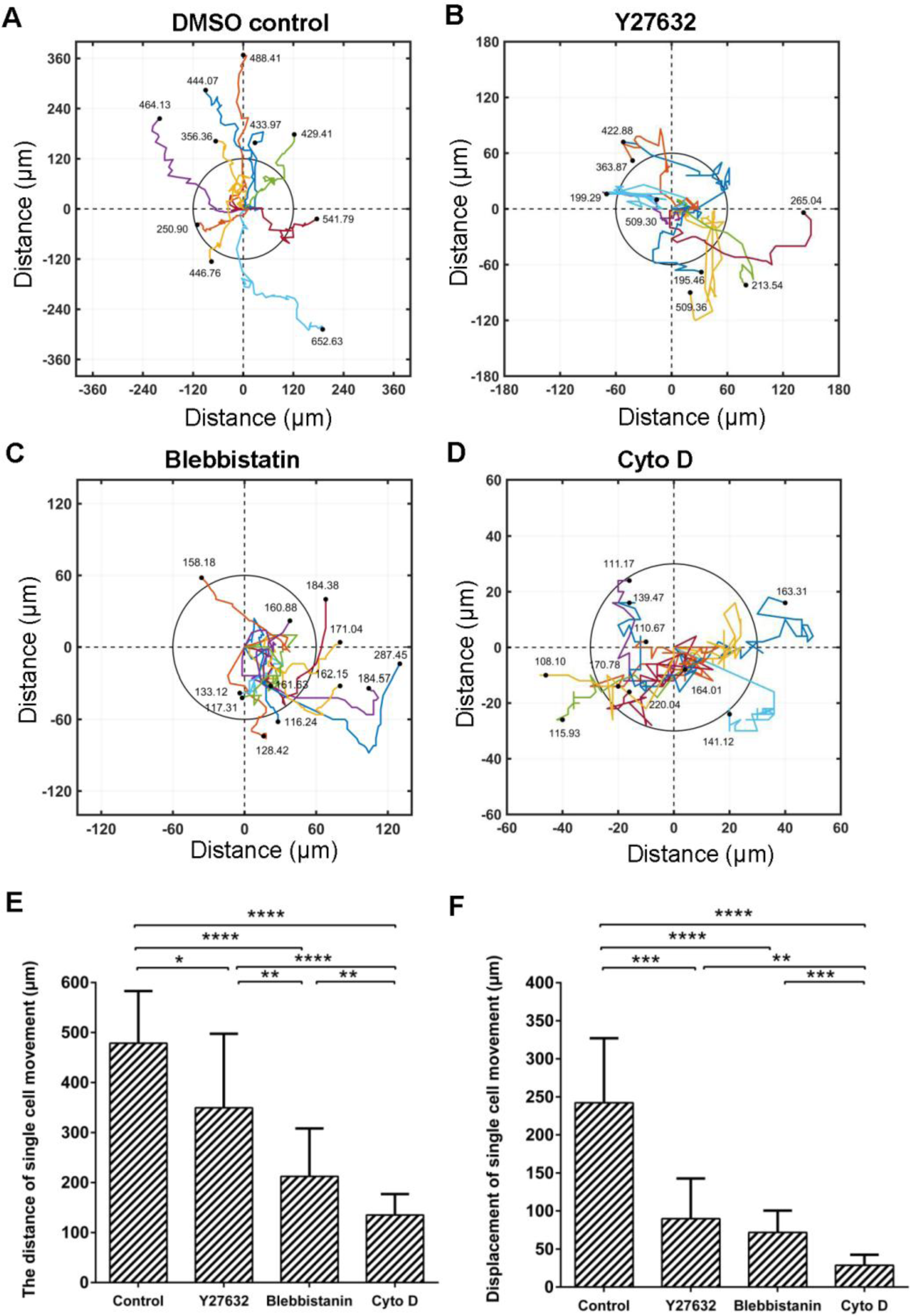
Tracking cell motions during the assembly of cell branching and COL fibers. Single cells in Figure 3 were analyzed for their translocations during the time courses of imaging. Their starting positions were normalized to zero, and the moving directions kept consistent with the cell migration directions on the images. (**A-D**) Tracks of the translocations of a group of single cells from the time courses under the control condition (DMSO) (A), with Y27632 (B), Blebbistatin (C), or Cyto D (D) treatment during the assembly of cell branching and COL fibers. (**E & F**) Quantifications and comparisons of the moving distances (Mean ± S.D.) (E) and final displacements (absolute values) (Mean ± S.D.) (F) from the original positions of those single cells under the indicated conditions. In statistical analysis, * represents P < 0.05, ** P < 0.01, *** P < 0.001, and **** P < 0.0001 for significant difference by Student’s t-test.

To further confirm whether cell motion mediated COL fiber assembly, we quantified the local cell movement and corresponding fiber fluorescence from the same group of time courses (from Fig. 2A(ii, iii) under the condition of contraction inhibition. Under the inhibition condition with Y27632 or Blebbistatin, further analysis found that those areas with relatively more active cell motions also generated stronger COL fibers, as shown in Fig. 4(A-C) in which there was positive correlationship between local cell mobility and fiber assembly from the same group of image samples. Taken together, these data suggest cell motions relying on contraction force and actin cytoskeleton facilitated COL fiber assembly from the medium during the branching formation.

**Figure 4.**
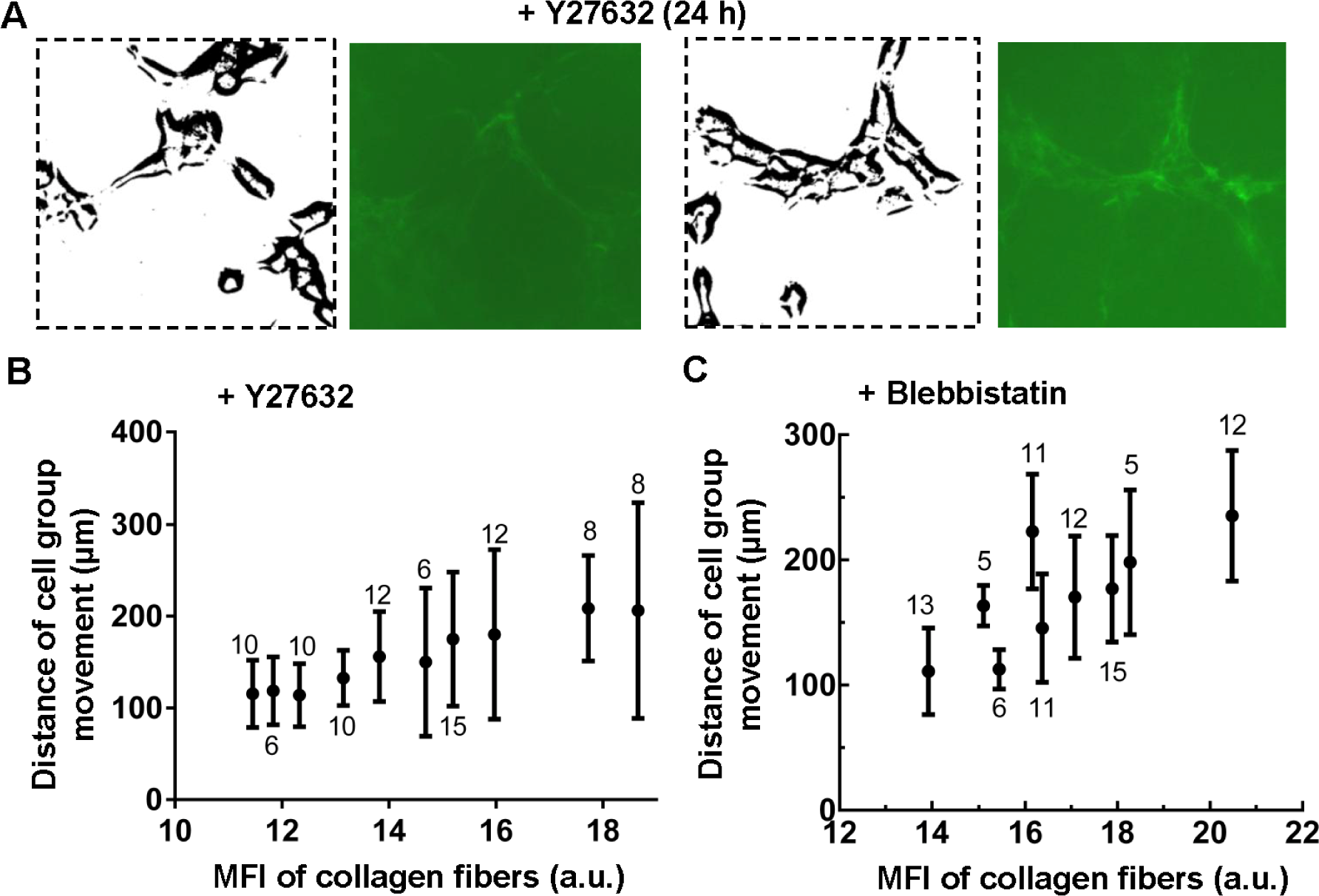
The positive correlation bewteen local cell movements and COL assembly under cell contraction inhibition. The local cell movements and COL fiber fluorescence were further analyzed based on the images of time courses from Figure 3 with Y27632 and Blebbistatin inhibitors. (**A**) The sample images with Y27632 treatment show the cells (binarized images) and relatively weaker or stronger fluorescent COL fibers. (**B&C**) The positive correlationship between local cell mobility and COL fiber assembly. From the same image group with 40 µM Y27632 (B) or 25 µM Blebbistatin (C) treatment, the mean cell moving distance and the mean COL fiber fluorescence at local regions were further quantified and plotted. The digitals beside the data indicate the cell numbers quantified at each region.

## Discussion

The components of ECM are secreted by cells *in vivo*, while ECM provides critical biochemical and mechanical communications with cells and tissues. The studies on matrix biology have been a long-time topic, and cells-assembled ECM represents more physiological structures and relevance [10,24,25]. In this work, we used fluorescent COL in the medium to exam how cells assembled the COL fibers on the top of 3-D Matrigel. This method provides a convenient way to visualize the dynamics of cell-based fiber assembly and to further investigate the underlying mechanism.

Our data showed that cell branching and COL fiber formation had a mutual-dependent relationship (Fig. 1), and the deposition of COL into the fibers was coordinated with the branching growth (Fig. 2). The recruitment of COL from the medium by the cells was an active process, which promoted branching formation. In considering that *in vivo* cells and tissues produce ECM components and help organize ECM physiological structures [10,13], our experiments provided evidence for the processes of cells-assisting COL fiber assembly, and cell branching morphogenesis shaped by COL fibers from *in vitro* study.

To investigate whether biomechanical force is involved into the fiber assembly, we demonstrated that inhibition of cell contraction force significantly reduced the strength of the fibers as measured by fluorescence intensity (Fig. 2A&C), and at the same time, cells only formed short and local branches (Fig. 2B). It is also worth noting that COL was still assembled into long but weaker fibers under the contraction inhibition conditions (Fig. 2A). When further looking at how intracellular contraction force regulated COL fiber assembly, tracking analysis showed cell motions, particularly directional migrations, were attenuated by inhibition of contraction force (Fig. 3A-C). Even in the same group of cells with contraction inhibition by Y27632 or Blebbistatin, those local regions with more cell motions had relatively stronger COL fibers (Fig. 4). Furthermore, disruption of cellular actin cytoskeleton almost blocked cell translocations, as well as completely inhibited both cell branching and COL fiber formations (Fig. 3A(iv)&C). So there was high coordination between both attenuated cell motions and fiber assembly, and both blocked cell translocation and fiber assembly. To put together, inhibition of cell contraction force reduced but didn’t block COL fiber assembly, while cell motion was closely relevant to the fiber growth.

COL fiber assembly/remodeling is highly regulated by biomechanical and biochemical processes [17,26,27]. ECM assembly and remodeling driven by cells represent more physiological processes occurring in vivo [9]. Our work revealed that cell mobility was an active factor in facilitating cells-mediated COL fibrillar assembly from the medium. This suggested new insights in matrix biology that cell motions might provide a biophysical microenvironment such as dynamics and strain/stress in assisting ECM assembly or remodeling. As a feedback, cells-assembled ECM structures actively joined in the developments of cell functions and morphogenesis such as cell branching formation here, indicating a precisely regulated biological system.

In summary, MDCK cells actively recruited COL molecules from the medium and regulated the fiber assembly, which in turn guided cell branching morphogenesis. The COL fiber assembly was mediated by cell contraction force and motions. We hypothesized that cell motions generated pulling force on COL fibers, which facilitated the fiber growth. Further studies will be needed to identify the molecular machinery of cells in recruiting and assembling COL fibers from the medium, and to discover the mechanism of possible strain-induced growth of fiber bundles.

## Acknowledgments

This work was supported financially by Natural Science Foundation of China (NSFC 11532003, 11872129, 31670950), Natural Science Foundation of Jiangsu Province (BK20181416), Jiangsu Provincial Department of Education, Natural Science Foundation of the Jiangsu Higher Education Institutions of China (18KJB180001), and Changzhou Science and Technology Bureau (CZSTB CZ20180017, CJ20190040).

## Author contributions

M. Ouyang and L. Deng designed the research; J. Wang performed the experiments and data analysis; J. Guo assisted in research design and experiments; M. Ouyang did data analysis & organization; B. Che quantified COL fiber fluorescence; L. Deng provided the setups of equipment; M. Ouyang, J. Wang, and L. Deng prepared the paper.

The authors declared that there is no competing interest in this work.

## References

[1] M. Affolter, R. Zeller, E. Caussinus, Tissue remodelling through branching morphogenesis, Nat Rev Mol Cell Biol 10 (2009) 831–842.

[2] M. Affolter, S. Bellusci, N. Itoh, B. Shilo, J.P. Thiery, Z. Werb, Tube or not tube: remodeling epithelial tissues by branching morphogenesis, Dev Cell 4 (2003) 11–18.

[3] T. Mammoto, A. Mammoto, D.E. Ingber, Mechanobiology and developmental control, Annu Rev Cell Dev Biol 29 (2013) 27–61.

[4] F. Kurth, K. Eyer, A. Franco-Obregon, P.S. Dittrich, A new mechanobiological era: microfluidic pathways to apply and sense forces at the cellular level, Curr Opin Chem Biol 16 (2012) 400–408.

[5] K.H. Vining, D.J. Mooney, Mechanical forces direct stem cell behaviour in development and regeneration, Nat Rev Mol Cell Biol 18 (2017) 728–742.

[6] B. Ladoux, R.M. Mege, Mechanobiology of collective cell behaviours, Nat Rev Mol Cell Biol 18 (2017) 743–757.

[7] C.L. Guo, M. Ouyang, J.Y. Yu, J. Maslov, A. Price, C.Y. Shen, Long-range mechanical force enables self-assembly of epithelial tubular patterns, Proc Natl Acad Sci U S A 109 (2012) 5576–5582.

[8] B.M. Baker, B. Trappmann, W.Y. Wang, M.S. Sakar, I.L. Kim, V.B. Shenoy, J.A. Burdick, C.S. Chen, Cell-mediated fibre recruitment drives extracellular matrix mechanosensing in engineered fibrillar microenvironments, Nat Mater 14 (2015) 1262–1268.

[9] A.D. Theocharis, D. Manou, N.K. Karamanos, The extracellular matrix as a multitasking player in disease, FEBS J 286 (2019) 2830–2869.

[10] G.A. Hoffmann, J.Y. Wong, M.L. Smith, On Force and Form: Mechano-Biochemical Regulation of Extracellular Matrix, Biochemistry (2019).

[11] A.J. Zollinger, M.L. Smith, Fibronectin, the extracellular glue, Matrix Biol 60-61 (2017) 27–37.

[12] K.C. Ingham, S.A. Brew, S. Huff, S.V. Litvinovich, Cryptic self-association sites in type III modules of fibronectin, J Biol Chem 272 (1997) 1718–1724.

[13] C. Zhong, M. Chrzanowska-Wodnicka, J. Brown, A. Shaub, A.M. Belkin, K. Burridge, Rho-mediated contractility exposes a cryptic site in fibronectin and induces fibronectin matrix assembly, J Cell Biol 141 (1998) 539–551.

[14] K.E. Kubow, R. Vukmirovic, L. Zhe, E. Klotzsch, M.L. Smith, D. Gourdon, S. Luna, V. Vogel, Mechanical forces regulate the interactions of fibronectin and collagen I in extracellular matrix, Nat Commun 6 (2015) 8026.

[15] Z. Kapacee, S.H. Richardson, Y. Lu, T. Starborg, D.F. Holmes, K. Baar, K.E. Kadler, Tension is required for fibripositor formation, Matrix Biol 27 (2008) 371–375.

[16] R. Zareian, K.P. Church, N. Saeidi, B.P. Flynn, J.W. Beale, J.W. Ruberti, Probing collagen/enzyme mechanochemistry in native tissue with dynamic, enzyme-induced creep, Langmuir 26 (2010) 9917–9926.

[17] B.P. Flynn, A.P. Bhole, N. Saeidi, M. Liles, C.A. Dimarzio, J.W. Ruberti, Mechanical strain stabilizes reconstituted collagen fibrils against enzymatic degradation by mammalian collagenase matrix metalloproteinase 8 (MMP-8), PLoS One 5 (2010) e12337.

[18] Z. Jia, T.D. Nguyen, A micromechanical model for the growth of collagenous tissues under mechanics-mediated collagen deposition and degradation, J Mech Behav Biomed Mater 98 (2019) 96–107.

[19] E.G. Canty, Y. Lu, R.S. Meadows, M.K. Shaw, D.F. Holmes, K.E. Kadler, Coalignment of plasma membrane channels and protrusions (fibripositors) specifies the parallelism of tendon, J Cell Biol 165 (2004) 553–563.

[20] S.J. Aper, A.C. van Spreeuwel, M.C. van Turnhout, A.J. van der Linden, P.A. Pieters, N.L. van der Zon, S.L. de la Rambelje, C.V. Bouten, M. Merkx, Colorful protein-based fluorescent probes for collagen imaging, PLoS One 9 (2014) e114983.

[21] M. Ouyang, S. Lu, Y. Wang, Genetically encoded fluorescent biosensors for live-cell imaging of MT1-MMP protease activity, Methods Mol Biol 1071 (2014) 163–174.

[22] M. Ouyang, R. Wan, Q. Qin, Q. Peng, P. Wang, J. Wu, M. Allen, Y. Shi, S. Laub, L. Deng, S. Lu, Y. Wang, Sensitive FRET Biosensor Reveals Fyn Kinase Regulation by Submembrane Localization, ACS Sens 4 (2019) 76–86.

[23] Q. Shi, R.P. Ghosh, H. Engelke, C.H. Rycroft, L. Cassereau, J.A. Sethian, V.M. Weaver, J.T. Liphardt, Rapid disorganization of mechanically interacting systems of mammary acini, Proc Natl Acad Sci U S A 111 (2014) 658–663.

[24] M.J. Lammi, J. Piltti, J. Prittinen, C. Qu, Challenges in Fabrication of Tissue-Engineered Cartilage with Correct Cellular Colonization and Extracellular Matrix Assembly, Int J Mol Sci 19 (2018).

[25] J.K. Mouw, G. Ou, V.M. Weaver, Extracellular matrix assembly: a multiscale deconstruction, Nat Rev Mol Cell Biol 15 (2014) 771–785.

[26] P. Lee, R. Lin, J. Moon, L.P. Lee, Microfluidic alignment of collagen fibers for in vitro cell culture, Biomed Microdevices 8 (2006) 35–41.

[27] N. Benseny-Cases, T.K. Karamanos, C.L. Hoop, J. Baum, S.E. Radford, Extracellular matrix components modulate different stages in beta2-microglobulin amyloid formation, J Biol Chem 294 (2019) 9392–9401.

